# The interspecific fungal hybrid *Verticillium longisporum* displays sub-genome-specific gene expression

**DOI:** 10.1101/341636

**Authors:** Jasper R.L. Depotter, Fabian van Beveren, Luis Rodriguez-Moreno, H. Martin Kramer, Edgar A. Chavarro Carrero, Gabriel L. Fiorin, Grardy C.M. van den Berg, Thomas A. Wood, Bart P.H.J. Thomma, Michael F. Seidl

## Abstract

Hybridization is an important evolutionary mechanism that can enable organisms to adapt to environmental challenges. It has previously been shown that the fungal allodiploid species *Verticillium longisporum*, causal agent of Verticillium stem striping in rape seed, has originated from at least three independent hybridization events between two haploid *Verticillium* species. To reveal the impact of genome duplication as a consequence of the hybridization, we studied the genome and transcriptome dynamics upon two independent *V. longisporum* hybridization events, represented by the hybrid lineages “A1/D1” and “A1/D3”. We show that the *V. longisporum* genomes are characterized by extensive chromosomal rearrangements, including between parental chromosomal sets. *V. longisporum* hybrids display signs of evolutionary dynamics that are typically associated with the aftermath of allodiploidization, such as haploidization and a more relaxed gene evolution. Expression patterns of the two sub-genomes within the two hybrid lineages are more similar than those of the shared A1 parent between the two lineages, showing that expression patterns of the parental genomes homogenized within a lineage. However, as genes that display differential parental expression *in planta* do not typically display the same pattern *in vitro*, we conclude that sub-genome-specific responses occur in both lineages. Overall, our study uncovers the genomic and transcriptomic plasticity during evolution of the filamentous fungal hybrid *V. longisporum* and illustrate its adaptive potential.

**Importance:** *Verticillium* is a genus of plant-associated fungi that include a handful of plant pathogens that collectively affect a wide range of hosts. On several occasions, haploid *Verticillium* species hybridized into the stable allodiploid species *Verticillium longisporum*, which is, in contrast to haploid *Verticillium* species, a Brassicaceae specialist. Here, we studied the evolutionary genome and transcriptome dynamics of *V. longisporum* and the impact of the hybridization. *V. longisporum* genomes display a mosaic structure due do genomic rearrangements between the parental chromosome sets. Similar to other allopolyploid hybrids, *V. longisporum* displays an ongoing loss of heterozygosity and a more relaxed gene evolution. Also, differential parental gene expression is observed, with an enrichment for genes that encode secreted proteins. Intriguingly, the majority of these genes displays sub-genome-specific responses under differential growth conditions. In conclusion, hybridization has incited the genomic and transcriptomic plasticity that enables adaptation to environmental changes in a parental allele-specific fashion.

## Introduction

Upon hybridization, two distinct genotypes are merged in a single organism. This surge in genomic variation can increase the adaptive potential of hybrid organisms, which may explain why stable hybrids are generally fitter than their parents in particular environments (1). However, hybrids may also encounter incompatibilities between parental genomes as they lack the recently shared evolutionary history (2). Hybridization can lead to the emergence of new species that are reproductively isolated from their parents, known as hybrid speciation (3, 4). Although the incidence of hybridization may be rare due to such incompatibilities, many organisms encountered hybridization at a particular point in their evolution (5). Hybridization has also impacted the evolution of humans, as our genomes still contain traces from Neanderthal introgression (6). Hybridization can occur between gametes after a conventional meiosis, leading to so-called homoploid hybrids. Alternatively, when compete sets of parental chromosomes combine, the hybridization is accompanied by genome duplication during so-called allopolyploidization.

Hybridization has impacted the evolution of a wide diversity of fungi (7–9). For instance, the yeast *Saccharomyces paradoxus*, a close relative of the baker’s yeast *Saccharomyces cerevisiae*, has naturally hybridized in North America forests (10), whereas also *S. cerevisiae* itself was shown to have undergone an ancient interspecies hybridization (11). Similarly, various *Candida* species that are opportunistic human pathogens display genomic traces of hybridization events (12–15). Hybridization also contributed to the evolution of various plant pathogenic fungi (7). Plant pathogens generally co-evolve with their hosts to evade host immunity, while hosts attempt to intercept pathogen ingress (16). In this process, plant pathogens secrete effector proteins that contribute to host immunity evasion and interfere with host metabolic processes (17), or affect to other processes to contribute to host colonization (18), such as the manipulation of host microbiomes (19, 20). Due to the increased adaptation potential, hybridization has been proposed as a potent driver in pathogen evolution as it can impact host interactions through increased virulence and host range alterations (8). For instance, the Ug99 strain of the wheat stem rust pathogen *Puccinia graminis* f. sp. *tritici* arose from a hybridization event and caused devastating epidemics in Africa and the Middle East (21, 22). Recent hybridization between the wheat powdery mildew, *B. graminis* f. sp. *tritici*, and rye powdery mildew, *B. graminis* f. sp. *secalis*, gave rise to the novel mildew species *Blumeria graminis* f. sp. *triticale* that, in contrast to its parents, is able to cause disease on triticale (23).

Upon hybridization, genomes typically experience a so-called “genome shock”, inciting major genomic reorganizations that can manifest by genome rearrangements, extensive gene loss, transposon activation, and alterations in gene expression (24, 25). Conceivably, these early stage alterations are primordial for hybrid survival, as divergent evolution is principally associated with incompatibilities between the parental genomes (26). Additionally, these initial re-organizations and further alterations in the aftermath of hybridization provide a source for environmental adaptation. Frequently, hybrid genomes lose their heterozygosity over time (27). Hybrids that are still sexually compatible with one of its parents can lose heterozygosity through backcrossing. Alternatively, heterozygosity can be a result of the direct loss of a homolog of one of the two parents (i.e. a homeolog) through deletion or through gene conversion whereby one of the copies substitutes its homeologous counterpart. Gene conversion and the homogenization of complete chromosomes played a pivotal role in the evolution of the osmotolerant yeast species *Pichia sorbitophila* (28). Two of its seven chromosome pairs consist of partly heterozygous and partly homozygous sections, whereas two chromosome pairs are completely homozygous. Gene conversion may eventually result in chromosomes consisting of sections of both parental origins, so called “mosaic genomes” (29). However, mosaic genomes can also arise through recombination between chromosomes of the different parents, such as in the hybrid yeast *Zygosaccharomyces parabailii* (30). Hybridization associated with polyploidy, allopolyploids can have an additional adaptive potential through the presence of an additional copy for most genes, which gives leeway to functional diversification (31, 32). Hybridization typically entails also alterations of gene expression patterns that are non-additive from the parental expression patterns (33, 34). Nevertheless, expression patterns are generally conserved upon hybridization, as the majority of allopolyploid genes are expressed in a similar fashion as their parental orthologs (35). For instance, more than half of the genes in an allopolyploid strain of the fungal grass endophyte *Epichloë* retained their parental gene expression pattern (36). Similar conservation has also been observed for *Blumeria graminis* f. sp. *triticale* as over half of the 5% most highly expressed genes are shared with both of its hybridization parents (37). In conclusion, the genomic and transcriptomic alterations accompanied with hybridization make that hybrids have a high potential for environmental adaptation (8).

Within the *Verticillium* genus that comprises nine haploid species, hybridization resulted in the emergence of the species *Verticillium longisporum* (38–41). *V. longisporum* is sub-divided into three lineages, each representing a separate hybridization event (39, 41). *Verticillium* species A1 is a parent of each of the three hybrids and hybridized with *Verticillium* species D1, D2 and D3, resulting in the *V. longisporum* lineages A1/D1, A1/D2, and A1/D3, respectively. Whereas species D2 and D3 have been classified as ‘likely *V. dahliae*’, species D1 has been classified as an enigmatic species that is closely related to *V. dahliae* (39). Species A1 is also an enigmatic species that diverged earlier from *V. dahliae* than the D1 species (39). Similar as the haploid *Verticillium* species, *V. longisporum* is thought to mainly undergo asexual reproduction, as a sexual cycle has never been described and populations are not outcrossing (40, 41). Interestingly, *V. longisporum* mainly infects plant hosts of the Brassicaceae family whereas other *Verticillium* species do not cause disease on Brassicaceous hosts (42). Moreover, while *V. dahliae* is characterized by an extremely broad host range that comprises hundreds of (non-Brassicaceae) plant species, *V. longisporum* only has a limited host range and hardly infects non-Brassicaceae species (42). After hybridization, *V. longisporum* conceivably encountered extensive genetic and transcriptomic alterations that facilitated its viability of a hybrid and the shift towards Brassicaceous hosts. In this study, we investigated the impact of allodiploidization on the evolution of *V. longisporum* by investigating genome, gene, and transcriptomic plasticity within and between two of the hybridization events.

## Results

### *Verticillium longisporum* displays a mosaic genome structure

The genomes of three *V. longisporum* strains from two different hybridization events were analyzed to investigate the impact of hybridization on genome structure. Previously, *V. longisporum* strains VLB2 and VL20, both belonging to the A1/D1 hybridization event, were sequenced with the PacBio RSII platform and assembled *de novo* (40). We now additionally sequenced the *V. longisporum* strain PD589, which originates from the A1/D3 hybridization event (39), using Oxford Nanopore sequencing technology and the BGISeq platform to obtain long reads and paired-end short reads, respectively. All *V. longisporum* genome assemblies were improved using chromatin conformation capture (Hi-C) sequencing that detects DNA-interactions (43). Moreover, centromeres can be located with Hi-C sequencing as they display strong interaction with centromeres in other chromosomes (44). We obtained genome assemblies of 72.7, 72.2 and 72.0 Mb consisting of 15, 15 and 16 pseudochromosomes for VLB2, VL20 and PD589, respectively (Figure 1A, Table 1). Every pseudochromosome contained a centromere, suggesting that the A1/D1 isolates have 15 chromosomes and that the A1/D3 isolate PD589 contains 16 chromosomes. However, chromosome 13 of strain PD589 displayed remarkably stronger DNA-interactions than the other chromosomes (Figure 1B, see green outline), as the median read coverage of chromosome 13 is 110x, whereas the read coverage is 58-70x for all other chromosomes (Figure S1). This finding suggests that chromosome 13 recently (partly) duplicated since the high sequence identity of the duplicated regions resulted in a collapsed assembly. Consequently, strain PD589 may therefore actually have 17 chromosomes in total.

**Figure 1.**
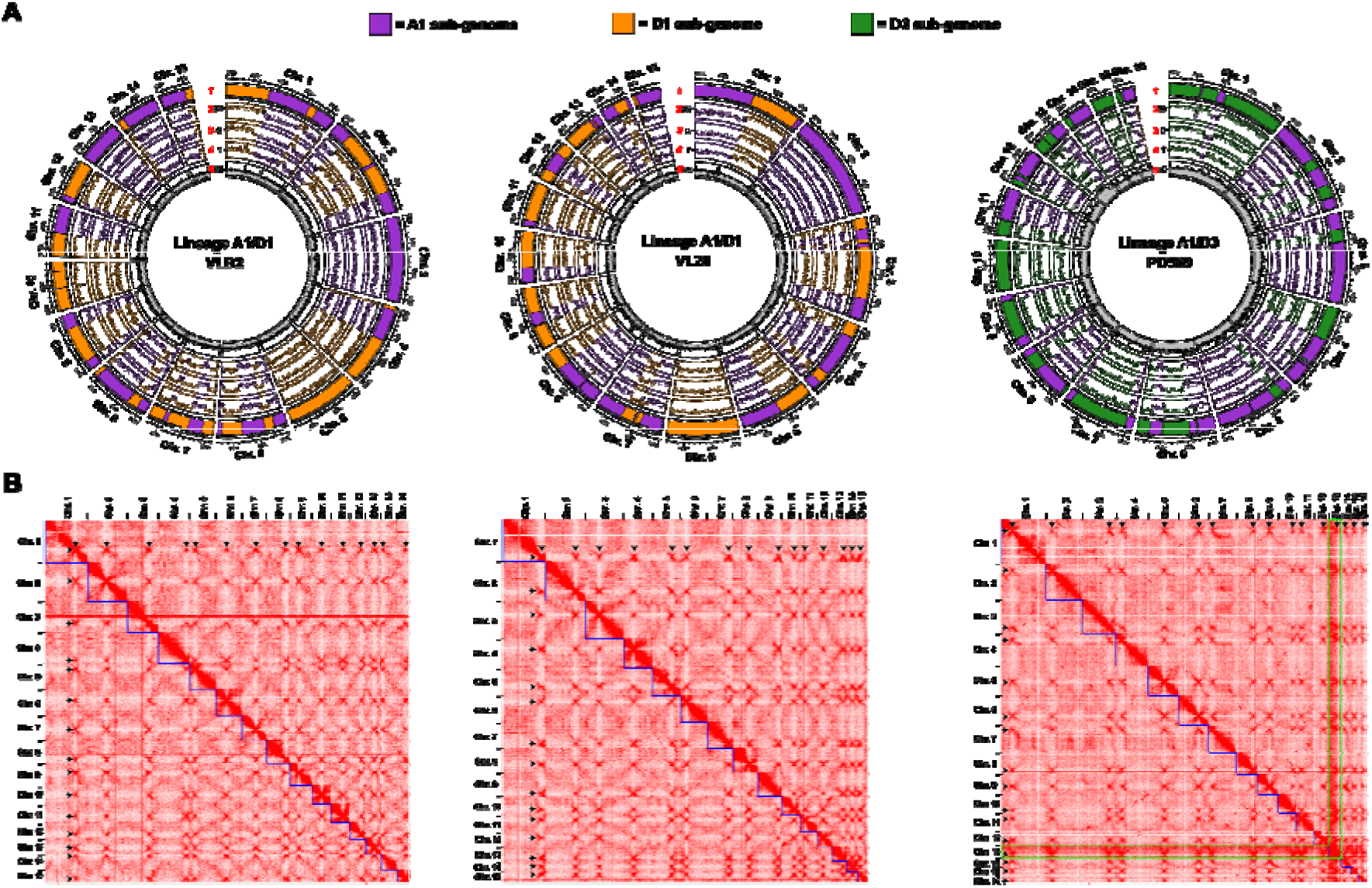
*Verticillium longisporum* displays a mosaic genome structure. (**A**) the *V. longisporum* chromosomes of strains VLB2, VL20 and PD589 are depicted. The different lanes in the circular plots represent: 1) regions assigned to Species A1 (purple), Species D1 (orange) and D3 (green); 2) sequence similarity of *V. longisporum* alignments to *V. dahliae* (% identity); 3) difference in sequence identity in percent point (pp) between exonic regions of *V. longisporum* double-copy genes. Only gene pairs with an ortholog in *V. dahliae* are depicted. Alleles with a higher identity to *V. dahliae* are depicted as a positive pp difference, whereas the corresponding homolog as a negative pp difference; 4) the relative difference in GC content (dGC) between genes in double copy; and 5) Read depth with non-overlapping windows of 10 kb. Data points of lanes 3-5 represent the average value of a window of eleven genes, which proceed with a step of one gene. (**B**) Hi-C contact frequency matrices for the three *V. longisporum* strains are shown. The red color indicates the contact intensity between genome region and the blue squares represent the pseudochromosomes. Centromeres display strong inter-chromosomal contacts and are visible as red dots outside the pseudochromosomes and are indicated with black arrows. Pseudochromosome 13 of strain PD589 generally displays stronger interactions than the other pseudochromosomes, which is outlined in green.

**Table 1.** Comparison of Verticillium longisporum and Verticillium dahliae genome assemblies.

|  | *V. longisporum* | *V. longisporum* | *V. longisporum* | *V. dahliae* |
| --- | --- | --- | --- | --- |
|  | VLB2<sup>1</sup> | VL20<sup>1</sup> | PD589 | JR2<sup>2</sup> |
| **Genome size** | 72.7 Mb | 72.2 Mb | 72.0 Mb | 36.2 Mb |
| **Assigned to A1 sub-genome** | 36.2 Mb | 36.5 Mb | 36.0 Mb | / |
| **Assigned to D sub-genome** | 35.9 Mb | 35.1 Mb | 34.7 Mb | / |
| **Undetermined** | 0.6 Mb | 0.6 Mb | 1.3 Mb | / |
| **Number of chromosomes** | 15 | 15 | 16/17<sup>4</sup> | 8 |
| **Number of predicted genes** | 18,679 | 18,592 | 18,251 | 9,636 |
| **Assigned to A1 sub-genome** | 9,342 | 9,343 | 8,961 | / |
| **Assigned to D sub-genome** | 9,298 | 9,188 | 9,229 | / |
| **Undetermined** | 39 | 61 | 61 | / |
| **Number of predicted genes encoding secreted proteins** | 2,084 | 2,049 | 1,960 | 1,071 |
| **Assigned to A1 sub-genome** | 1,052 | 1,041 | 952 | / |
| **Assigned to D sub-genome** | 1,025 | 1,004 | 1000 | / |
| **Undetermined** | 7 | 4 | 8 | / |
| **Repeat content** | 14.55% | 14.54% | 12.78% | 11.69% |
| **BUSCO completeness<sup>3</sup>** | 99.1% | 99.3% | 97.9% | 98.6% |

<sup>1</sup>Previously published assemblies were reassembled using Hi-C sequencing (40)
<sup>2</sup>(46)
<sup>3</sup>Based on Ascomycota Benchmarking Universal Single Copy Orthologs (BUSCOs)
<sup>4</sup>Total chromosome number is uncertain as PD589 contains one (partially) duplicated chromosome

Being able to determine the parental origin of individual genomic regions is elementary to investigate genome evolution in the aftermath of hybridization. As the D parents of *V. longisporum* hybridizations (D1 and D3) are phylogenetically closer related to *V. dahliae* than parent A1 (39), *V. longisporum* genome alignments to *V. dahliae* display a bimodal distribution with minima around 96.0% identity (Figure S2). To separate the two sub-genomes, we assigned regions with an average sequence identity to *V. dahliae* smaller than this minimum to parent A1 whereas regions with an identity larger than this threshold were assigned to the D parent (Figure 1A). In this manner, 36.0-36.5 Mb were assigned to the A1 parents and 34.7-35.9 Mb to the D parents (Table 1). Thus, the sub-genome sizes are quite similar for each of the isolates and correspond to the expected genome sizes of haploid *Verticillium* species (44, 45). Intriguingly, the majority of the *V. longisporum* chromosomes is composed of DNA regions that originate from different parents, and only two chromosomes have a single parental origin in each of the strains (Figure 1, Table S1). Thus, *V. longisporum* chromosomes generally are mosaics of DNA regions of different parental origin.

### The mitochondrial genome is inherited from the A1 parent in all lineages

To determine the phylogenetic position of the parental sub-genomes of the *V. longisporum* hybridization, we used the *V. longisporum* sub-genome sequences and previously published genome sequences of the haploid *Verticillium* species (45, 46) to construct a phylogenetic tree based on 1,520 Ascomycete benchmarking universal single copy orthologs (BUSCOs) that were present in a single copy in all analyzed *Verticillium* lineages. In accordance with previous phylogenetic studies (39, 40), the A1 parents diverged earlier from *V. dahliae* than the D1 and D3 parents (Figure S3). Furthermore, the D1 parent diverged earlier from *V. dahliae* than the D3 parent. We also constructed a phylogenetic tree based on mitochondrial DNA to determine the parental origin on the mitochondria. The *V. longisporum* mitochondrial genomes were assembled in a single contig with overlapping ends, indicating their circular nature. The mitochondrial genomes of the three *V. longisporum* strains were all 26.2 kb in size and were more than 99.9% identical in sequence. The phylogenetic position of the *V. longisporum* mitochondrial genomes clusters with the mitochondrial genomes of *V. alfalfae* and *V. nonalfalfae* (Figure S3). As the mitochondrial genome sequence is almost identical for three strains that are derived from the two hybridization events, the common A1 parent is the likely donor of the mitochondria.

### Genomic rearrangements are responsible for the mosaic nuclear genome

Typically, a mosaic structure of a hybrid nuclear genome can originate from gene conversion or from chromosomal rearrangements between DNA strands of different parental origin (27). To analyze the extent of gene conversion, protein-coding genes were predicted for the *V. longisporum* strains using BRAKER with RNA-Seq data from fungal cultures grown *in vitro* (47). The number of predicted genes ranged from 18,251 to 18,679 for the different *V. longisporum* strains, which is 89-94% higher than the genes number of *V. dahliae* strain JR2 predicted using the same methodology (9,636 genes) (Table 1). In total, 8,961-9,343 genes were assigned to the sub-genome of parent A1, whereas the number of genes in the D3 sub-genomes ranged from 9,188 to 9,298 (Table 1). Thus, the gene numbers are similar for the different *V. longisporum* sub-genomes and comparable to the gene number of *V. dahliae*. Over 79% of the *V. longisporum* genes have one homolog, i.e. they occur in two copies, which can originate from gene duplication (paralogy) or from the hybridization event (homeology) (Figure 2). Within each of the *V. longisporum* sub-genomes, most genes (96.9-99.6%) have no additional homolog and occur in a single copy (Figure 2B), indicating that most homologous gene pairs in each *V. longisporum* genome are homeolog in nature and that gene conversion only played a minor role after hybridization. To find traces of gene conversion during their evolution, the sequence identity of 6,213 genes that have two homologous copies in the two A1/D1 strains was compared, as these two strains belong to distinct populations (40). Only eight genes were found to be highly similar (<1% nucleotide sequence diversity) in VLB2, whereas the corresponding gene pair in VL20 was more diverse (>1%) (Figure 3A). Similarly, in *V. longisporum* strain VL20, six highly similar copies were found that are more divergent in VLB2, thereby confirming that gene conversion has hitherto only played a marginal role in the evolutionary aftermath of the *V. longisporum* hybridization.

**Figure 2.**
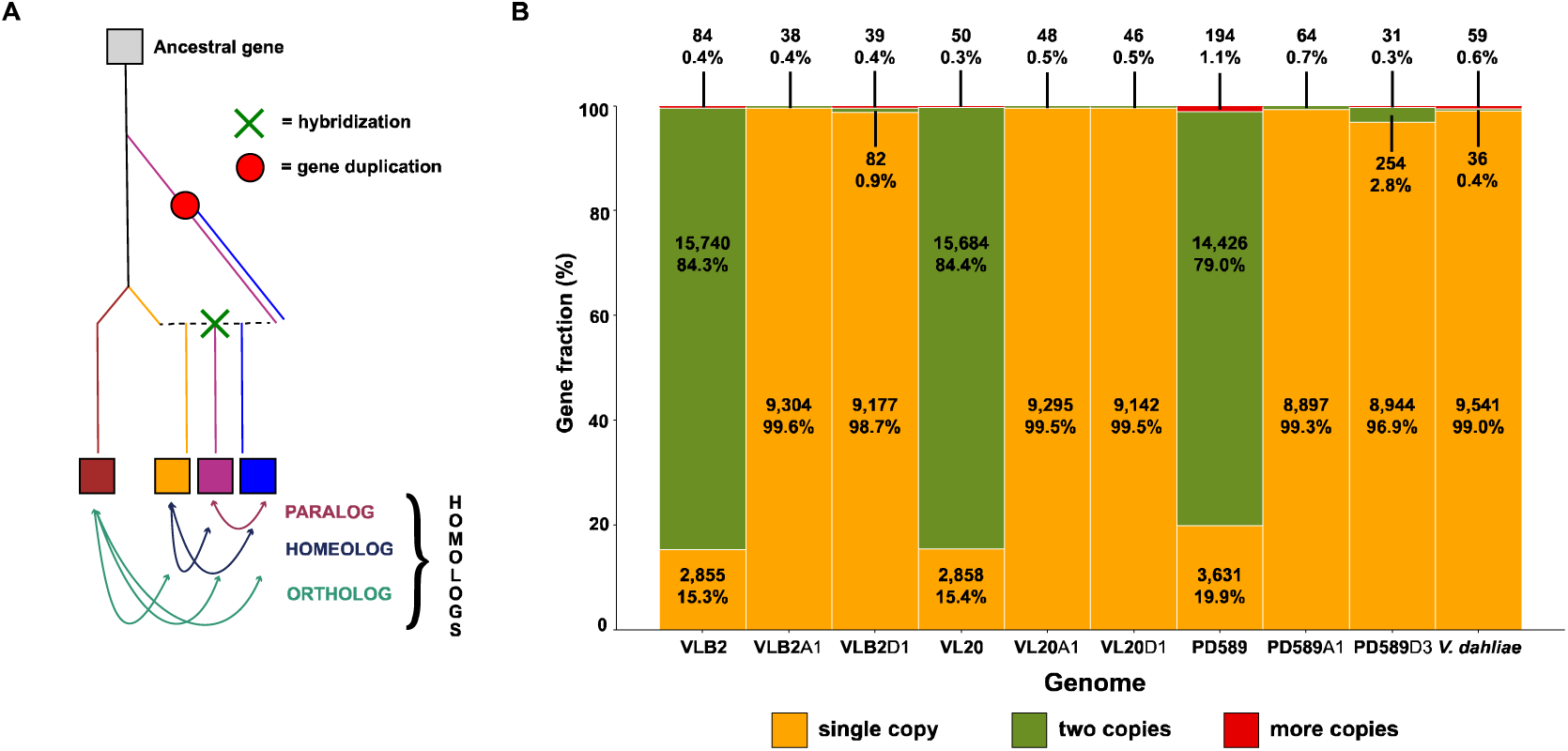
*V. longisporum* genes with two copies are almost exclusively homeologs. (**A**) Schematic overview of different evolutionary origins of homologous genes in hybrids. Paralogs are homologous genes that originate from gene duplication, while orthologous genes originate by speciation. Homeologs are homologous genes originating from a hybridization event. (**B**) The gene fraction occurring in single, two, and more than two copies in the *V. longisporum* strains VLB2, VL20 and PD589 are shown, with *V. dahliae* (strain JR2) as comparison. “A1”, “D1” and “D3” represent species A1, D1 and D3 sub-genomes, respectively.

**Figure 3.**
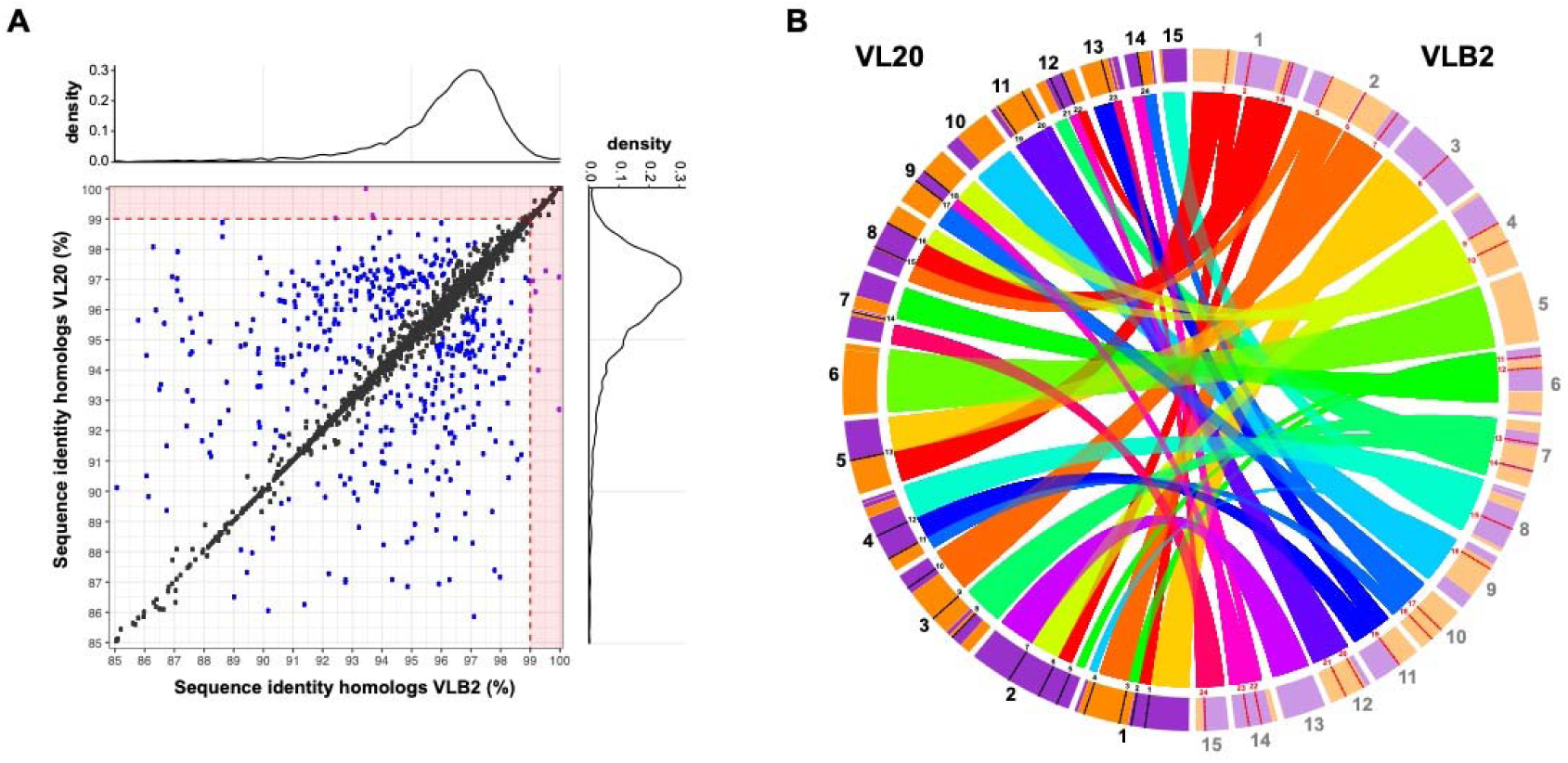
The mosaic genome structure of *Verticillium longisporum* originates from genomic rearrangements. (**A**) The contribution of gene conversion to *V. longisporum* genome evolution. Sequence identities between genes in copy, present in *V. longisporum* VLB2 and VL20, are depicted. Homologous gene pairs within a strain that encountered gene conversion are expected to have higher similarity within a strain compared with the corresponding gene pair in the other strain. Gene pairs with divergence of more than one percent in one *V. longisporum* strain and less than one percent in the other strain were considered conserved in the latter strain (purple dots in the red zones). In other cases, pairs that differ less than one percent are depicted as a black dot, whereas a difference greater than one percent is depicted as a blue dot. (**B**) The contribution of genomic rearrangements to *V. longisporum* genome evolution. The *V. longisporum* chromosomes of strains VLB2 (displayed on the right) and VL20 (displayed on the left) are depicted. Ribbons indicate syntenic genome regions between the two strains and contig colors indicate the parental origin similar to Figure 1 (purple = A1 and orange = D1). The red and black lines with the associated numbers on the chromosomes indicate synteny breaks.

Considering that gene conversion played only a minor role during genome evolution (Figure 3), the mosaic genome structure of *V. longisporum* likely originated from rearrangements between homeologous chromosomes. To identify chromosomal rearrangements after the hybridization event that led to the A1/D1 lineage, the genome of *V. longisporum* strain VLB2 was aligned to that of strain VL20, revealing 24 syntenic breaks (Figure 3B). Rearrangement occurred in the majority of the chromosomes as only 2 and 3 chromosomes did not have syntenic breaks in VLB2 and VL20, respectively (Figure 3B). As genomic rearrangements are often associated with repeat-rich genome regions, such as in *V. dahliae* (46, 48, 49), the synteny break points were tested for their association to repetitive regions. Since the median repeat fraction in a 20 kb window around the repeats is 15.5%, which is significantly more than the median repeat fraction based on random sampling (average = 3.4%, σ = 1.7%) (Figure S4), it can be concluded that the chromosomal rearrangements are similarly associated with repeats also in *V. longisporum*. In conclusion, chromosomal rearrangement rather than gene conversion is the main mechanism explaining the mosaic structure of the *V. longisporum* genome.

### *V. longisporum* loses heterozygosity through deletions

To study putative gene losses in the aftermath of hybridization, we determined genes that have no homeolog or paralog, and can thus be considered to occur in single copy. For the A1/D1 isolates, 15.3-15.4% of the genes occur in single copy, whereas this is 19.9% for A1/D3 isolate PD589 (Figure 2B). We checked if proteins encoded by single copy genes are enriched for particular Gene Ontology (GO) terms, Clusters of Orthologous Groups (COGs), or encoding a protein with a signal peptide, which suggests that these proteins are secreted. No GO terms and COGs were enriched for the single copy genes in any of the *V. longisporum* strains (Fisher’s exact test with Benjamini-Hochberg correction, *p*-value < 0.05). In total, 7.8-10.2% of the single copy genes encodes a protein with a signal peptide, which is a significantly lower than the 11.9-12.3% for genes with a homologous copy in the same genome (Fisher’s exact test, *p*-value < 0.05). Of the A1/D1 single copy genes, 52% reside in the A1 sub-genome and 47% in the D1 sub-genome. Similarly, for PD589, 49% and 50% reside in the A1 and D3 sub-genome, respectively. Thus, single copy genes are equally distributed across the two sub-genomes in *V. longisporum*. Single copy genes can either originate from gene loss or from parent-specific contributions to the hybrid. Since VLB2 and VL20 originate from the same hybridization event (40), we can quantify how many single copy genes originate from gene loss during divergence of VLB2 and VL20. In total, 14.7-14.8% of the singly copy genes have at least one copy in each sub-genome of the other A1/D1 strain, suggesting that gene deletion is an on-going process in *V. longisporum* evolution. Of the single copy genes that lost their homeolog after the hybridization event, 48% resided in the species A1 sub-genome, whereas 51-52% in the D1 sub-genome, suggesting that gene losses occurred to a similar extent in each of the sub-genomes.

### Acceleration of gene evolution upon hybridization

To investigate the gene sequence evolution subsequent to hybridization, we compared the ratio of non-synonymous (*Ka*) and synonymous (*Ks*) substitutions (ω) for branches leading to *Verticillium* species (Figure 4). To exclude the putative impact of the (partial) chromosome 13 duplication in PD589, we excluded genes of this chromosome from the analysis. Substitution rates were determined for a total of 3,823 genes that have just one ortholog in the analyzed *Verticillium* species, *V. alfalfae*, *V dahliae*, *V. nonalfalfae* and *V. nubilum*, as well as in each of the *V. longisporum* sub-genomes. To mitigate possible biases of different divergence times between the *Verticillium* species, we performed the analyses four times: three times with the two sub-genomes of *V. longisporum* strains VLB2, VL20, and PD589, and once with *V. dahliae* and the A1 sub-genome of VLB2 (Figure 4). *V. longisporum* and *V. dahliae* genes with higher ω than their *V. alfalfae*, *V. nonalfalfae* and *V. nubilum* orthologs were considered quickly evolving, whereas those with lower ω were considered slowly evolving. Comparing the D1/D3/*V. dahliae* branch, *V. dahliae* has 839 slowly evolving genes, which is a higher number than the 758 and 629 slowly evolving genes of the *V. longisporum* D1 and D3 sub-genomes, respectively. Conversely, *V. dahliae* has 1,229 quickly evolving genes, which is lower than the number found for the *V. longisporum* D1 and D3 sub-genomes, 1,357/1,372 (VL20/VLB2) and 1,586, respectively (Figure 4). This observation fits to the prevailing hypothesis that hybridization accompanied by genome duplication has a ‘relaxing’ effect on gene evolution (32, 50). Furthermore, the lower number of slowly evolving genes and larger number of quickly evolving genes in the D3 sub-genome is significantly different from the D1 sub-genome (Fisher’s exact test, *P* < 0.001). Similar to the D sub-genomes, the A1 sub-genome of lineage A1/D3 has higher number of quickly evolving genes (2,072 vs. 1,691-1,714) and lower number of slowly evolving genes (462 vs. 628-634) than the A1 sub-genome of lineage A1/D1. In conclusion, *V. longisporum* lineage A1/D3 genes generally evolve faster than lineage A1/D1 genes in both sub-genomes. This may indicate that A1/D3 evolved a longer time under the more relaxes gene evolutionary conditions than A1/D1, i.e. A1 and D3 hybridized a longer time ago than A1/D1.

**Figure 4.**
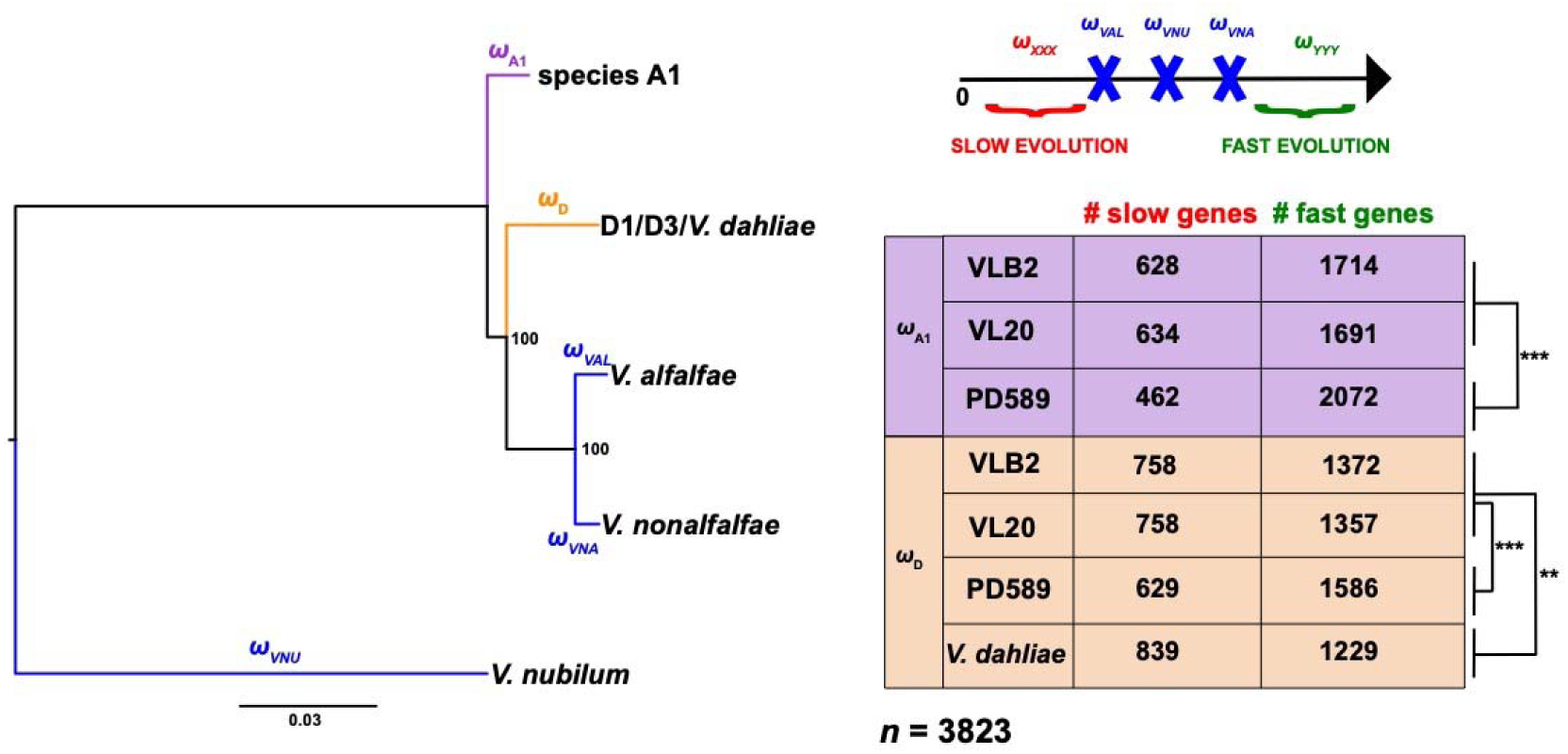
*Verticillium longisporum* genes divergence faster than *Verticillium dahliae* orthologs. *Ka/Ks* ratios (ω) were calculated for the tree branches leading to *Verticillium* spp. of the clade Flavnonexudans genomes and the *V. longisporum* sub-genomes. A total of 3,823 genes with one ortholog in all respective *Verticillium* (sub-)genomes were analyzed. *V. longisporum* and *V. dahliae* genes with fast or slow evolution have a higher ω or lower ω, respectively, than their *V. alfalfae*, *V. nonalfalfae* and *V. nubilum* orthologs. Significance in gene numbers was calculated with the Fisher’s exact test. **: *P* < 0.01 and ***: *P* < 0.001.

To see whether particular genes evolve faster, we functionally characterized the *V. longisporum* A1/D3 genes that have a higher ω than their *V. alfalfae*, *V. nonalfalfae* and *V. nubilum* orthologs, but also higher than their lineage A1/D1 homologs from the corresponding A1 and D sub-genomes to select genes that quickly evolved after the A1 and D1/D3 last common ancestor. In total, 1,350 of the 3,823 (35.3%) analyzed genes were quickly evolving in PD589 A1 sub-genome and 1,084 (28.4%) for the D3 sub-genome. We screened for GO term, COG and secreted protein enrichments in these fast evolving A1/D3 genes and no enrichments for the COGs and for genes encoding secreted proteins were found.

In the A1 sub-genome 3 GO terms with a molecular function were significantly enriched, associated with molecule binding (protein and ATP) and ATPase activity. In the D3 sub-genome, “ATP binding” was the only significantly enriched GO term, which was also enriched in the A1 sub-genome. In conclusion, the more pronounced “gene relaxation” in the A1/D3 lineage when compared with the A1/D1 lineage does not clearly seem to affect genes with particular functions.

### Expression pattern homogenization in the hybridization aftermath

To investigate the impact of hybridization on gene expression, the expression of *V. longisporum* genes was compared with *V. dahliae* orthologs from strains grown *in vitro* in potato dextrose broth. To this end, expression of single copy *V. dahliae* genes was compared with orthologs that are present in two homeologous copies in three *V. longisporum* strains (VLB2, VL20, and PD589). Genes on chromosome 13 from strain PD589 and their homologs were excluded from the analysis to avoid putative biases due to a (partial) chromosome duplication, and in total 5,604 expressed genes were compared. RNA sequencing reads were mapped to the predicted *V. longisporum* genes of which 50-51% mapped to species A1 homeologs and 49-50% to the D homeologs. Thus, we observed no global differences in overall contribution to gene expression of the sub-genomes. Over half of the *V. longisporum* homeologs display no differential expression with their *V. dahliae* ortholog, indicating that the majority of the genes did not evolve differential expression patterns (Figure 5A). In both lineages, higher numbers of differently expressed genes were found in the A1 sub-genome than in the D sub-genomes; 27 vs. 23% for A1/D1 and 38 vs. 34% for A1/D3, respectively. The higher fraction of differentially expressed A1 genes is in accordance with the more distant phylogenetic relationship of parent A1 with *V. dahliae* than of the D parents (Figure S3). Intriguingly, although D3 diverged more recently from *V. dahliae* than D1, D3 has more differentially expressed orthologs with *V. dahliae* than D1. When comparing expression patterns between sub-genomes, 11-13% of the genes display differential expression between their A1 and D homeologs. Intriguingly, this is more than half the number of differentially expressed D and *V. dahliae* orthologs (23-34%), despite the fact that the D parents diverged more recently from *V. dahliae* than from species A1 (Figure S3). In general, the gene expression patterns of the A1 and D sub-genomes of the same hybridization event are highly correlated (0.93-0.96), higher than D sub-genomes and *V. dahliae* strain JR2 (0.85-0.89) and higher than the A1 sub-genomes between hybridization events (0.82-0.84) (Figure 5B; Table S2). To compare these expression patterns with the gene expression variation between different *V. dahliae* strains, we sequenced RNA from the cotton-infecting *V. dahliae* strain CQ2 grown in potato dextrose broth. Although JR2 and CQ2 belong to the same species, their overall gene expression pattern is more dissimilar (ρ = 0.89) than that of *V. longisporum* sub-genomes (Figure 5B; Table S2). The overall discrepancy in phylogenetic relationship and expression pattern similarities suggests that sub-genome expression patterns of the sub-genomes in *V. longisporum* homogenized upon hybridization.

**Figure 5.**
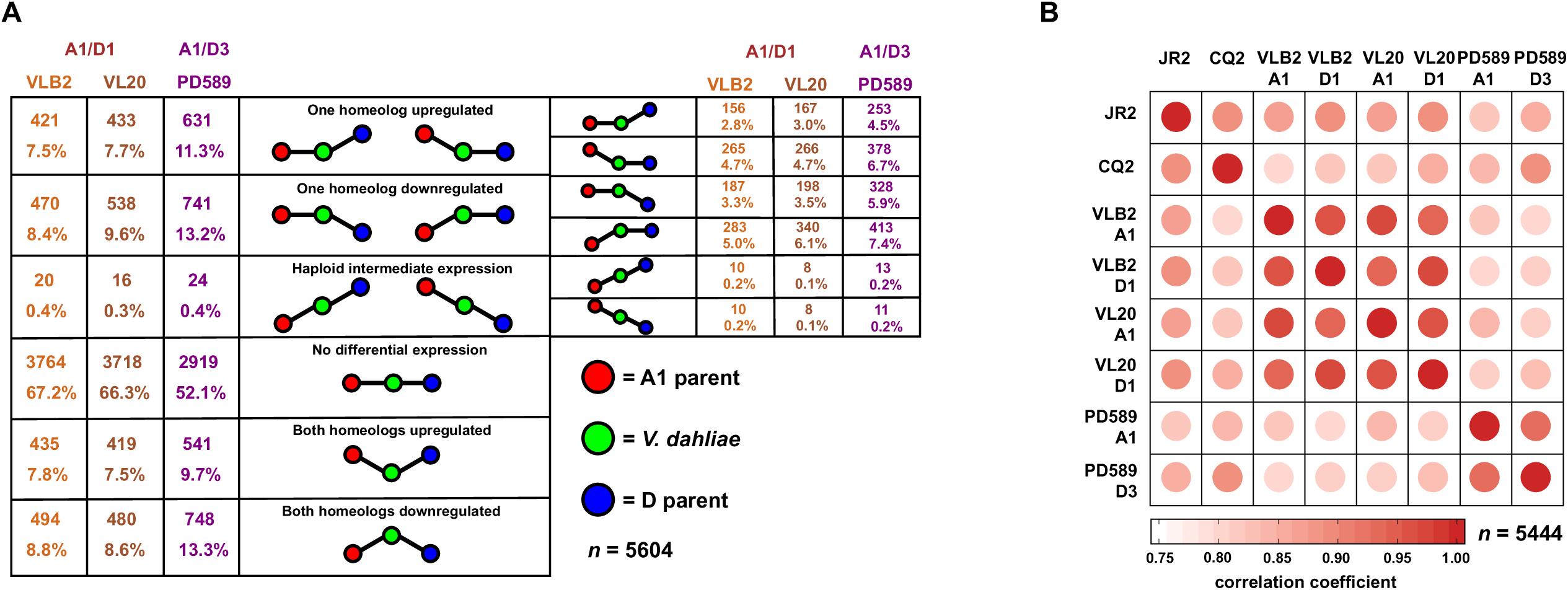
Gene expression patterns of *Verticillium longisporum* sub-genomes display remarkable resemblance. Expression pattern comparison between *Verticillium longisporum* sub-genomes and *Verticillium dahliae* in culture medium (**A**) Differential expression between *V. longisporum* and *V. dahliae* genes. Only genes with one homolog in *V. dahliae* and two homeologs in the *V. longisporum* strains VLB2, VL20 and PD589 were considered for differential expression. The significance of differential expression was calculated using *t*-tests relative to a threshold of log2-fold-change of 1 and a Benjamini-Hochberg corrected *p*-value cut-off of 0.05. (**B**) Expression pattern correlation between *V. longisporum* and *V. dahliae*. Only genes with one homolog in *V. dahliae* strains JR2 and CQ2 and two homeologs in the *V. longisporum* strains VLB2, VL20 and PD589 were considered. Spearman’s correlation coefficients (ρ) were calculated based on the mean transcripts per million values of three replicates.

### Differential homeolog expression occurs in particular gene categories

Although parental gene expression patterns appear to have globally homogenized upon hybridization, differential homeolog expression occurs as well (Figure 5). To assess if genes with differential homeolog expression belong to specific gene groups, we screened for functional enrichments. In total, 10% of the fast-evolving PD589 genes (defined in the previous section) have different homeolog expression, which is significantly lower than the 12% different homeolog expression for the remainder of the genes (Fisher’s exact test, *P* < 0.05). In both A1/D1 and A1/D3 lineages, genes with differential homeolog expression are enriched for GO terms related to oxidation-reduction processes, transmembrane transport and FAD binding (Figure 6A, Table S3). Additionally, the COGs “carbohydrate transport and metabolism” and “secondary metabolites biosynthesis, transport, and catabolism” (Q) are enriched in both lineages (Table S3). Furthermore, we tested if genes encoding secreted proteins were significantly enriched among the genes with differential homeolog expression. Indeed, 23 and 16 % of the genes with different homeolog expression code for a secreted protein in the lineage A1/D1 isolates and in the A1/D3 isolate, respectively, whereas this is 9% of the genes that do not display differential expression among homeologs (VLB2 *P* =1.23E-32, VL20 *P* =3.71E-29 and PD589 *P* =1.14E-08, Fisher’s exact test). In conclusion, differential homeolog expression seems to be important for particular gene categories, including categories that can be implicated in plant pathogenicity.

**Figure 6.**
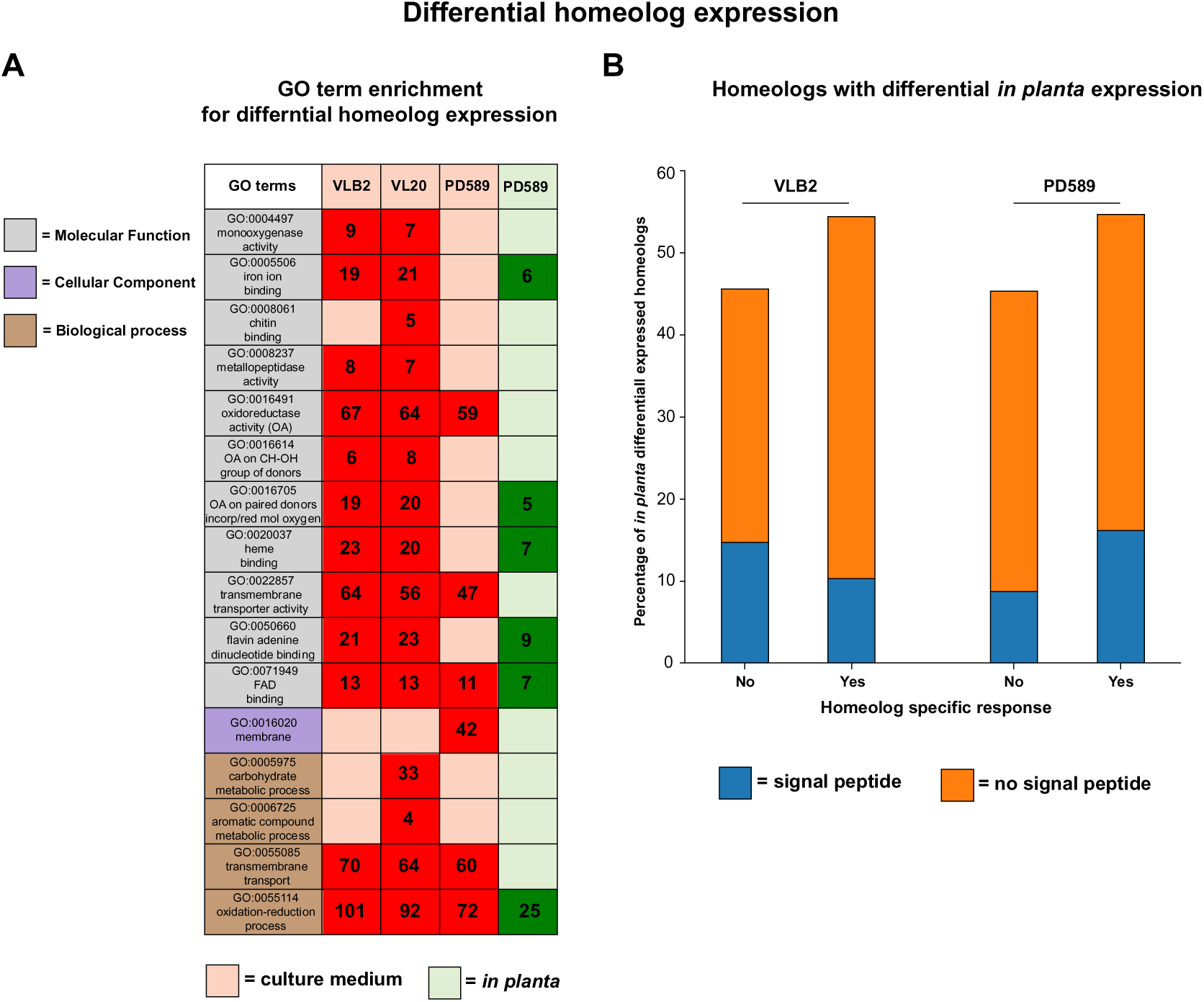
*Verticillium longisporum* displays sub-genome-specific gene expression responses. Functional enrichments for *Verticillium longisporum* genes with different homeolog expression in culture medium and *in planta*. Only *V. longisporum* genes with two homeologs were considered. (**A**) Gene Ontology (GO) terms that are significantly enriched in differentially expressed homeologs of VLB2, VL20 and PD589 are displayed. A more detailed overview and level of significance are reported in Table S3. The number of genes with differential homeolog expression are indicated. (**B**) Fractions of genes with differential homeolog expression *in planta* with and without a homeolog specific response. Genes have a homeolog specific response as they display differential homeolog expression *in planta* and have no differential or the opposite expression ratio for *V. longisporum* grown in culture medium.

### Homeolog-specific expression responses upon plant colonization

Considering the plant pathogenic nature of *V. longisporum*, and also that genes encoding secreted proteins, which are often implicated in pathogenicity on host plants, are enriched among the genes with differential homeolog expression, we assessed homeolog-specific gene expression during plant colonization. To this end, oilseed rape plants were inoculated with the *V. longisporum* strains VLB2, VL20, and PD589. As observed previously, oilseed rape plants inoculated with VLB2 and PD589 developed typical *Verticillium* symptoms including stunted plant growth and leaf chlorosis (51). In contrast, oilseed rape plants inoculated with VL20 did not display any disease symptoms. Consequently, we performed total RNA sequencing for oilseed rape plants inoculated with *V. longisporum* strains VLB2 and PD589. For strain PD589, genes on chromosome 13 and their homeologs were removed from the analysis. For VLB2 and PD589, 51% of the reads mapped to the A1 sub-genome and 49 to the D sub-genome. Thus, similar to *in vitro* grown *V. longisporum*, we did not observe any global difference in overall contribution to gene expression of one of the sub-genomes *in planta*. In total, 1.1% and 2.7% of the homeologs displayed differential expression *in planta*, which is less than the 11.3 and 13.4% found for VLB2 and PD589 grown *in vitro*, respectively. Genes with differential homeolog expression *in planta* were not enriched for any GO term in the A1/D1 strain VLB2 (Table S3), whereas in the A1/D3 strain PD589, differentially expressed homeologs were enriched for GO terms associated with oxidation-reduction processes and molecular binding (iron ion, heme, flavin adenine dinucleotide and FAD) (Figure 6A, Table S3). For A1/D1 and A1/D3, genes with different homeolog expression were enriched for those encoding secreted proteins; 25% of differentially expressed homeologs encode secreted proteins and 8-9% of the non-differentially expressed homeologs encode other proteins (*P* < 0.05, Fisher’s exact test). Thus, similar to *in vitro V. longisporum* grown, differential homeolog expression *in planta* is especially important for genes encoding secreted proteins. In 33% of these secretome genes with differential homeolog expression *in planta*, no Pfam domain could be annotated, which is a feature often observed for effector proteins as they are often examples of biological innovation (52). Of these genes that could be functionally annotated, a carbohydrate-active enzyme (CAZyme) function was annotated in 32% of the cases. The remaining part of the functionally annotated genes with differential homeolog expression included other enzymes, such as proteases, lipases, carboxylesterases and peroxidases. We compared genes with differential homeolog expression *in planta* and *in vitro* to assess potential correlation. Intriguingly, over half (54-55%) of the differentially expressed homeologs *in planta* are not differentially expressed in culture medium or have the inverse expression pattern, e.g. *in vitro*: A1>D and *in planta* A1<D (Figure 6B). Thus, over half of the genes with a differential homeolog expression *in planta* display a homeolog-specific response compared to *in vitro* growth. For VLB2, 19% of these genes with a homeolog-specific response encode secreted proteins, whereas 32% of genes with similar differential homeolog expression *in planta* and *in vitro* encode secreted proteins. The opposite pattern in observed for PD589, i.e. 30% with a homeolog-specific response and 19% with similar differential homeolog expression *in planta* and *in vitro*. However, these differences were not significant (*P* > 0.05, Fisher’s exact test). In conclusion, different growing conditions cause homeolog-specific changes in the majority of the *V. longisporum* genes with differential homeolog expression, which is enriched in genes that encode secreted proteins.

## Discussion

Hybridization is a powerful evolutionary mechanism that can lead to the emergence of new plant pathogens with distinct features when compared with their parents (8, 23). Here, we reveal the transcriptomic plasticity of the hybrid pathogen *V. longisporum* and illustrate the parental allele-specific response to different environmental cues. Differential expressed *V. longisporum* homeologs are enriched for genes encoding secreted proteins that generally act to facilitate environmental manipulation (53). Interestingly, over half of the differentially homeolog expressed genes *in planta* display different relative contributions *in vitro*. Thus, upon the environmental changes that are associated with different growth conditions, *V. longisporum* encounters sub-genome specific gene expression alterations, leading to differential homeolog expression. Although not previously reported for any other hybrid plant pathogen, sub-genome specific gene expression alterations has previously been reported to occur in the artificial yeast hybrid *S. cerevisiae* x *Saccharomyces uvarum* upon temperature change (54). Genes with these sub-genome specific responses were involved in a variety of biological processes, including the trehalose metabolic process that is involved in thermotolerance. Thus, more generally, hybrid fungi, comprising natural as well as artificial hybrids, respond to environmental change in an allele-specific manner, especially for genes that manipulate or mitigate environmental changes. Secretome genes with a differential homeolog expression *in planta* often have an enzymatic function or lack an annotated Pfam domain, which is a feature often observed for effector proteins that act in pathogenicity (52). Thus, conceivably, homeolog specific responses *in planta* occurs in genes that are important for host colonization. Similarly, differential homeolog expression in the hybrid opportunistic human pathogen *Candida orthopsilosis* involves genes that are implicated in host interactions, related to superoxide dismutase activity and zinc metabolism (55).

Although differential homeolog expression occurs, the general tendency is that expression patterns between the A1 and D sub-genomes homogenized upon hybridization (Figure 5). Despite the absence of A1 and D1 species due to their enigmatic nature, we can conclude that parental gene expression patterns homogenized in the aftermath of hybridization as sub-genome expression patterns display more resemblance than the expression pattern between *V. dahliae* and the D parents and between the A1 sub-genomes of different hybridization events (Figure 5B; Table S2). Homogenization of parental expression patterns has been similarly observed in the fungal allopolyploid *Epichloë* Lp1 (36) as well as in the artificial hybrid *S. cerevisiae* x *S. uvarum* where the extent of differential ortholog expression between the parents was diminished upon hybridization (56). Thus, gene expression homogenization seems to be a more general phenomenon in fungi. Gene expression divergences may evolve through mutations in regulatory sequences of the gene itself (*cis*-effects), such as promoter elements, or alterations in other regulatory factors (*trans*-effects), such as chromatin regulation (57, 58). Conceivably, the higher correlation in homeolog expression patterns than parental ortholog expression patterns originates from changes in *trans* regulators as homeologs, in contrast to orthologs, share the same nuclear environment (58). Intriguingly, parent D3 has more genes that are differentially expressed to *V. dahliae* orthologs than parent D1, despite that D3 diverged more recently from *V. dahliae* than D1 (Figure 5, Figure S4). Correspondingly, the A1 sub-genome of the lineage A1/D3 displays more differential gene expression with *V. dahliae* than the A1 sub-genome of the A1/D1 lineage. This can indicate that A1 and D3 hybridized before A1 and D1, as more distinct expression patterns may have evolved over time.

In addition to the transcriptomic plasticity of homeolog expression upon environmental changes, *V. longisporum* is also plastic on a genomic level, which is displayed by its mosaic structure (Figure 1A, Table S1). Mosaicism is also observed in the grass pathogen *Zymoseptoria pseudotritici*, which is a close relative of the prominent wheat pathogen *Zymoseptoria tritici* (29). *Z. pseudotritici* is a homoploid hybrid that displays mosaicism on a population level where genome regions inherited from one parent display low variation, whereas high variable genome regions were transmitted from both parents. *V. longisporum* mosaicism is caused by extensive genomic rearrangements after hybridization (Figure 2B, 3). Genomic rearrangements are major drivers of evolution and facilitate adaptation to novel or changing environments (48). Genomic rearrangements are not specific to the hybrid nature of *V. longisporum* as other *Verticillium* species similarly encountered extensive chromosomal reshuffling (44, 45, 49, 59). In *V. dahliae*, genomic rearrangements especially occur in genomic regions that were originally described as lineage-specific regions, which are enriched for active transposable elements, and that are derived from segmental duplications that were followed by extensive reciprocal gene losses, encounter nucleotide sequence conservation and have an unique epigenomic profile (49, 59–62). These lineage-specific regions are enriched for *in planta* expressed genes and contain effector genes that facilitate host infection (59, 60, 63, 64). Since more recently, these lineage-specific regions are referred to as dynamic chromosomal regions (60). Similar to *V. dahliae*, syntenic breaks in *V. longisporum* often reside in repeat-rich genome regions as repetitive sequences (Figure S3), due to their abundance, are more likely to act as a substrate for unfaithful repair of double-strand DNA breaks (48, 49). However, the presence of two genomes within a single hybrid nucleus may also provide homeologous sequences with sufficient identity to mediate unfaithful repair.

The *V. longisporum* D genomes globally display accelerated evolution when compared with their *V. dahliae* orthologs (Figure 4), which may be a consequence of genome doubling. Interestingly, the *V. longisporum* A1/D3 lineage strain PD589 encountered a more divergent gene evolution when compared with the A1/D1 lineage strains VLB2 and VL20 in both sub-genomes, indicating that the A1/D3 hybridization occurred prior to the A1/D1 hybridization as a longer allodiploid state could facilitate extended sequence divergence (65). However, accelerated evolution is not consistently observed in fungi as deceleration upon allopolyploidization has been recorded in the fungal genus *Trichosporon* (66). Arguably, environmental cues play an important role in the speed and grade of gene diversification upon allopolyploidization (67). Possibly, accelerated gene evolution in *V. longisporum* is cued by a host range alteration as it is, in contrast to haploid *Verticillium* species, a Brassicaceae specialist (42). However, we did not find functional enrichments in fast evolving genes that points towards that hypothesis. Moreover, as the A1 species remains enigmatic, we cannot be sure a host shift occurred (39, 41).

Whole-genome duplication events are typically followed by extensive gene loss, often leading to reversion to the original ploidy state (68). For instance, the artificial interspecific hybrid *S. cerevisiae* x *S. uvarum* encountered nine independent events where loss of heterozygosity occurred after evolving for hundreds of generations under nutrient-limited conditions (69). Heterozygosity loss has only proceeded to a limited extent in *V. longisporum*, as 84% of lineage A1/D1 genes and 79% of lineage A1/D3 genes are present in two copies, whereas the haploid *V. dahliae* only contains 0.4% of its genes in two copies (Figure 2B). Thus, the *V. longisporum* genome displays the symptoms of a recent allodiploid, with gene loss being an on-going process that by now has only progressed marginally. Heterozygosity loss can indicate deleterious epistatic interactions between parental genomes that need to homogenize in order for the hybrid to be viable. Similar to other fungal hybrids (69, 70), we did not observe a specific group of genes where loss of heterozygosity was selected for. The degree of haploidization is a third indication that the A1/D3 lineage likely hybridized prior to A1/D1, as haploidization progress further in A1/D3 than in A1/D1 (Figure 2B). *C. orthopsilosis* hybrids from different hybridization events have different degrees of heterozygosity loss but genes were homeologs are maintained in both hybrids are enriched for those to have differential homeolog expression (55). Although species often revert to their original ploidy state after polyploidization, a retention of both homeolog copies can also be evolutionary advantageous, for instance to respond in a parental allele-specific fashion to environmental cues (Figure 6).

## Conclusion

Allodiploidization is an intrusive evolutionary mechanism in fungi where two chromosome sets from parents with a distinct evolutionary history merge. Consequently, most genes obtain an additional gene copy that can be differentially regulated according to the environmental conditions. Thus, allodiploid fungi can respond in a parental allele-specific fashion to environmental cues. Besides such parental allele-specific gene expression, allodiploidization furthermore contributed to a dynamic genome evolution through rearrangements between parental chromosome sets and accelerated gene evolution in *V. longisporum*. Thus, in comparison to haploid *Verticillium* species, *V. longisporum* has a high adaptive potential that can contribute to host immunity evasion and may explain its specialization towards Brassicaceous plant hosts.

## Material and Methods

### *V. longisporum* genome sequencing and assembly

Genome assemblies of the *V. longisporum* strains VLB2 and VL20 were previously constructed using long reads obtained through single-molecule real-time (SMRT) sequencing (40). Here, we sequenced *V. longisporum* strains PD589 using Oxford Nanopore Technology (ONT). In order to obtain DNA of PD589, spores were harvest from PDA plates and grown in 1/5 PDB for 5 days. Mycelium and spores were collected on Myra cloth, freeze-dried overnight and ground to fine powder. For DNA isolation, 100 mg of material was used and incubated for one hour at 65°C with 800 μL DNA extraction buffer (0.35 M Sorbitol, 0.1 M Tris-base, 5 mM EDTA pH 7.5), nucleic lysis buffer (0.2 M Tris, 0.05 M EDTA, 2 M NaCl, 2% CTAB) and Sarkosyl (10% w/v) in a 2:2:1 ratio. Subsequently, ½ volume of phenol/chloroform/isoamyl alcohol (25:24:1) was added, shaken vigorously and incubated at room temperature (RT) for 5 minutes before centrifugation at maximum speed (16,000 rpm) for 15 minutes (RT). The upper (aqueous phase) layer was transferred to a new tube, 5 μL of RNAase (10 mg/μL) was added and incubated at 37°C for one hour. Next, ½ volume of chloroform was added, mixed and centrifuged at maximum speed for 10 minutes at RT. The upper layer was transferred to a new tube and a second chloroform wash step was performed. After transferring the upper layer to a new tube, it was mixed with 1 volume (∼ 800 μL) of 100% ice-cold ethanol by gently inverting the tube and finally the DNA was fished out and washed twice by applying 500 μL of 70% ethanol. Finally, the DNA was air-dried, resuspended in nuclease-free water and stored at 4°C overnight. The DNA quality, size and quantity were assessed by nanodrop, gel electrophoresis and Qubit analyses, respectively.

To sequence the *V. longisporum* strain PD589 DNA, a library was prepared as described in the manufactures protocol provided by ONT (SQK-RAD004) with an initial amount of ∼ 400 ng HMW DNA. The library was loaded onto a R9.4.1 flow cell which ran for 24 hours and yielded ∼7 Gb of data. ONT sequencing reads were basecalled using Guppy (version 3.1.5) using the high accuracy base calling algorithm. Subsequently, adapter sequences were identified and removed using Porechop (version 0.2.3; default settings); adapters at the end of the reads were trimmed and reads with internal adapters were discarded. To be able to polish the genome assembly, we used the same HWA DNA isolated for ONT sequencing to generate ∼35 million high-quality (95% > phred score of 20) 150 bp paired-end reads (∼76x coverage) using the BGISeq platform (BGI Tech Solutions, Hongkong).

The *V. longisporum* PD589 genome was *de novo* assembled using Canu (version 1.8; genomeSize=70m, corOutCoverage=100, batOptions=’-dg 3 -db 3 -dr 1 -ca 500 -cp 50’) (71). In total, 924,740 cleaned ONT reads were used for the *de novo* assembly of which 743,753 where >1 kb (∼88x coverage). The genome assembly was polished using two sequential rounds of Apollo (version 1.1) (72). To this end, the high-quality paired-end reads were mapped to the genome assembly using bwa (version 0.7.17-r1188; default settings) (73).

To improve the assemblies to (near) chromosome level, chromatin conformation capture (Hi-C) followed by high-throughput sequencing was performed for VLB2, VL20 and PD589, similar as previously reported (44). For the three *V. longisporum* strains, one million spores were added to 400 ml Potato Dextrose Broth and incubated for 6 days at 22°C with continuous shaking at 120 rpm. 300 mg (fresh weight) mycelium was used as input for generating Hi-C sequencing libraries with the Proximo Hi-C kit (Microbe) (Phase Genomics, Seattle, WA, USA), according to manufacturer instructions. Hi-C sequencing libraries were paired-end (2×150 bp) sequenced on the NextSeq500 platform at USEQ (Utrecht, the Netherlands). Juicer (v1.6) was then used to map Hi-C sequencing reads to the previously obtained assemblies (74). The contact matrices generated by Juicer were used by the 3D *de novo* assembly (3D-DNA) pipeline (v180922) to eliminate misjoints in the previous assemblies (75). The assemblies were manually further improved using Juicebox Assembly Tools (JBAT) (v.1.11.08) (76). JBAT was subsequently used to determine centromere location based on intra- and inter-chromosomal contact frequencies. Only contigs that were larger than 100 kb were maintained in the assembly. Coverage of the ONT for the *V. longisporum* PD589 assembly was determined for 20 kb windows with samtools depth (v1.9) (77) and reads were mapped with minimap2 (v2.17-r941) (78).

The mitochondrial genomes of the haploid *Verticillium* species were previously sequenced and assembled (45). Mitochondrial *V. longisporum* genomes were assembled alongside the nuclear genomes (40). Mitochondrial contigs consisted of multiple copies of the mitochondrial genome due to its circular nature. A single copy of the mitochondrial genome was excised using BEDTools getfasta (v2.23.0) (Quinlan and Hall 2010). Filtered *V. longisporum* sub-reads were mapped to these single-copy mitochondrial assemblies using circlator (v1.5.5) (Hunt et al. 2015). The mapped reads were subsequently used to make a new *V. longisporum* mitochondrial genome assembly using SAMtools mpileup (v1.8) (Li et al. 2009).

### RNA sequencing

To obtain RNA-seq data for *Verticillium* grown in culture medium, *V. dahliae* isolates JR2 and CQ2, and *V. longisporum* isolates VLB2, VL20 and PD589 were grown for three days in potato dextrose broth (PDB) with three biological replicates for each isolate. To obtain RNA-seq data from *in planta* growth, two-week-old plants of the susceptible oilseed rape cultivar ‘Quartz’ were inoculated by dipping the roots for 10 minutes in 1×10^6^ conidiospores ml^-1^ spore suspension of *V. longisporum* isolates VLB2, VL20 and PD589, respectively (51). After root inoculation, plants were grown in individual pots in a greenhouse under a cycle of 16 h of light and 8 h of darkness, with temperatures maintained between 20°C and 22°C during the day and a minimum of 15°C overnight. Three pooled samples (10 plants per sample) of stem fragments (3 cm) were used for total RNA extraction. Total RNA was extracted based on TRIzol RNA extraction (Simms et al. 1993). cDNA synthesis, library preparation (TruSeq RNA-Seq short-insert library), and Illumina sequencing (single-end 50 bp) was performed at the Beijing Genome Institute (BGI, Hong Kong, China).

### Gene prediction and functional characterization

The *V. longisporum* assemblies of strains VLB2, VL20 and PD589 and the previously published assemblies of *V. dahliae* strains JR2 and CQ2 (46, 61) were annotated using the BRAKER v2.1.4 pipeline with RNA-Seq data with the options “--softmasking” and “-- fungus” enabled (47). RNA-seq reads from *Verticillium* grown in axenic culture (all replicates) were mapped to the assemblies using TopHat v2.1.1 (79). Predicted genes with internal stop codons, without a start codon or with an unknown amino acid in the encoded protein sequence were removed from the analysis. The secretome prediction was done using SingalP5 (v5.0) (80). Pfam and Gene Ontology (GO) function domains were predicted using InterProScan (v5.42-78.0) (81). Clusters of Orthologous Group (COG) categories were determined for protein sequences using eggNOG-mapper (v2.0) with the taxonomic scope set on Ascomycota (82, 83). Carbohydrate-Active enzymes (CAZymes) were annotated using the dbCAN2 meta server (84, 85). A protein was considered a CAZyme if at least two of the three tools (HMMER,DIAMOND and Hotpep) predicted a CAZyme function.

### Parental origin determination

Sub-genomes were divided based on the differences in sequence identities between species A1 and D1/D3 with *V. dahliae*. *V. longisporum* genomes of VLB2, VL20 and PD589 were aligned to the complete genome assembly of *V. dahliae* JR2 using NUCmer (v 3.1), which is part of the MUMmer package (86). Here, only 1-to-1 alignments longer than 10 kb and with a minimum of 80% identity were retained. Subsequent alignments were concatenated if they aligned to the same contig with the same orientation and order as the reference genome. The average nucleotide identity was determined for every concatenated alignment and used to divide the genomes into sub-genomes. Differences in GC-content between homologous genes present in two copies were calculated as described (28). GC content of gene coding regions were calculated with infoseq from EMBOSS (v.6.6.0.0) (87). The features to indicate the biparental origin of the *V. longisporum* genomes were visualized using the R package circlize (v.0.4.10) (88).

### Genome analysis

The quality of genome assemblies was assessed by screening the presences of Benchmarking Universal Single-Copy Orthologs (BUSCOs) using the BUSCO software version 4.0.6 with the database “ascomycota_odb10” (89).

Repeats were *de novo* identified using RepeatModeler (v1.0.11) and combined with the repeat library from RepBase (release 20170127) (90). The genomic location of repeats was identified with RepeatMasker (v4.0.6).

The phylogenetic relationship of the nuclear and mitochondrial (sub-)genomes of the *Verticillium* species of the clade Flavnonexudans (38), using following haploid strains: *V. alfalfae* = PD683, *V. dahliae* = JR2, *V. nonalfalfae* = TAB2 and *V. nubilum* = PD621 (45, 46). Phylogenetic trees based on nuclear DNA were constructed based on the Ascomycete BUSCOs that were shared by all the included species (89). Nucleotide sequences were separately aligned using MAFFT (v7.464) (91). Phylogenetic trees were inferred using RAxML with the GTRGAMMA substitution model (v8.2.11) (92). The robustness of the inferred phylogeny was assessed by 100 rapid bootstrap approximations.

Homologs in *Verticillium* were determined using nucleotide BLAST (v2.2.31+). Genes with a minimum identity of 80% and a minimum overlap of 80% were considered homologs, which was determined using the SiLiX (v.1.2.10-p1) software (93).

Global nucleotide alignments using the Needle-Wunsch algorithm of the EMBOSS package were used to determine homologous gene pairs in VLB2 and VL20 (v6.6.0.0) (87). Sequence identity between these genes in copy were determined based on their global alignment. Synteny between the VLB2 and VL20 genome assemblies was determined by using one-to-one alignments obtained by NUCmer (v 3.1), which is part of the MUMmer package (86). The synteny was visualized with the R package circlize (v.0.4.10) (88).

### Gene divergence

Previously published annotations of the haploid *Verticillium* species *V. dahliae*, *V. alfalfae*, *V. nonalfalfae*, *V. nubilum*, *V. tricorpus* and *V. albo-atrum* were used to compare the evolutionary speed of orthologs (45, 46). The VESPA (v1.0b) software was used to automate this process (94). The coding sequences for each *Verticillium* species were filtered and subsequently translated using the VESPA ‘clean’ and ‘translate’ function. Homologous genes were retrieved by protein BLAST (v2.2.31+) querying a database consisting of all *Verticillium* protein sequences. Here, the options “-max_hsps 1” and “-qcov_hsp_perc 80” were used. Homologous genes were grouped with the VESPA ‘best_reciprocal_group’ function. Only homology groups that comprised a single representative for every *Verticillium* spp. were used for further analysis. Protein sequences of each homology group were aligned with muscle (v3.8) (95). The aligned protein sequences of the homology groups were conversed to nucleotide sequence by the VESPA ‘map_alignments’ function. The alignments were used to calculate *Ka/Ks* for every branch of the species phylogeny using codeml module of PAML (v4.9) with the following parameters: F3X4 codon frequency model, wag.dat empirical amino acid substitution model and no molecular clock (96). To this end, this phylogenetic tree topology was used: ((((*V. dahliae*/D1/D3,(*V. alfalfae*, *V. nonalfalfae*)),A1),*V. nubilum*),(*V. tricorpus*, *V. albo-atrum*)). Divergence was only compared for genes that are present in the two sub-genomes of the *V. longisporum* strains VLB2, VL20 and PD589.

### Gene expression analysis

The RNA sequencing reads were filtered using the Trinity software (v2.9.1) option trimmomatic under the standard settings (97). The reads were then mapped to the *Verticillium* genomes using Bowtie 2 (v2.3.5.1) with the first 15 nucleotides on the 5’-end of the reads being trimmed because of inferior quality (98). To compare gene expression patterns, homologs were retrieved by nucleotide blast BLAST (v2.2.31+). Genes with a minimum identity of 80% and a minimum overlap of 80% were considered homologs, which was determined using the SiLiX (v.1.2.10-p1) software (93). Reads were counted to the predicted gene coding regions using the R package Rsubread (v1.34.7) Significant differential expression of a locus was calculated using the R package edgeR (v3.26.8) (99). Significance of differential expression was calculated using t-tests relative to a threshold of log2 fold change of 1 with Benjamini-Hochberg correction using a *p*-value cut-off of 0.05.

### Data accession

Raw RNAseq reads and genome assemblies are deposited at NCBI under the BioProject PRJNA473305.

## Acknowledgements

The authors would like to thank the Marie Curie Actions program of the European Commission that financially supported the research of J.R.L.D. Work in the laboratories of B.P.H.J.T. and M.F.S is supported by the Research Council Earth and Life Sciences (ALW) of the Netherlands Organization of Scientific Research (NWO). B.P.H.J.T acknowledges support from the Deutsche Forschungsgemeinschaft (DFG, German Research Foundation) under Germanýs Excellence Strategy – EXC 2048/1 – Project ID: 390686111. The funders had no role in study design, data collection and analysis, decision to publish, or preparation of the manuscript. We thank Sander Y.A. Rodenburg for sharing bioinformatics scripts.

