## Supplemental data for "The interspecific fungal hybrid *Verticillium longisporum* displays sub-genome-specific gene expression"

**Table S1. Fractions of individual *Verticillium longisporum* chromosomes that belong to the A1 and D parent.**

| **Chr.** | **VLB2** | | | **VL20** | | | **PD589** | | |
| --- | --- | --- | --- | --- | --- | --- | --- | --- | --- |
|  | **A1** | **D1** | **UND** | **A1** | **D1** | **UND** | **A1** | **D3** | **UND** |
| **1** | 54% | 45% | 1% | 55% | 44% | 1% | 8% | 92% | 0% |
| **2** | 39% | 60% | 1% | 100% | 0% | 0% | 75% | 23% | 2% |
| **3** | 99% | 0% | 1% | 35% | 64% | 0% | 85% | 15% | 1% |
| **4** | 39% | 60% | 1% | 67% | 31% | 2% | 54% | 43% | 3% |
| **5** | 0% | 99% | 0% | 53% | 47% | 0% | 85% | 14% | 1% |
| **6** | 56% | 43% | 1% | 0% | 99% | 1% | 46% | 50% | 4% |
| **7** | 28% | 71% | 0% | 72% | 25% | 2% | 17% | 81% | 1% |
| **8** | 76% | 22% | 2% | 69% | 30% | 1% | 68% | 31% | 1% |
| **9** | 39% | 61% | 0% | 20% | 80% | 0% | 27% | 72% | 1% |
| **10** | 0% | 100% | 0% | 30% | 70% | 1% | 0% | 99% | 1% |
| **11** | 56% | 42% | 2% | 13% | 86% | 1% | 65% | 29% | 6% |
| **12** | 13% | 86% | 1% | 47% | 52% | 1% | 79% | 19% | 2% |
| **13** | 98% | 2% | 1% | 23% | 77% | 1% | 41% | 59% | 0% |
| **14** | 87% | 13% | 1% | 55% | 44% | 2% | 81% | 18% | 2% |
| **15** | 79% | 20% | 1% | 91% | 8% | 1% | 0% | 100% | 0% |
| **16** |  |  |  |  |  |  | 72% | 23% | 5% |

**Table S2. Expression pattern correlation of genes between *Verticillium dahliae* and *Verticillium longisporum* sub-genomes grown in culture medium.**

|  | **JR2** | **CQ2** | **VLB2 A1** | **VLB2 D1** | **VL20 A1** | **VL20 D1** | **PD589 A1** | **PD589 D3** |
| --- | --- | --- | --- | --- | --- | --- | --- | --- |
| **JR2** | 1.00 | 0.89 | 0.87 | 0.89 | 0.87 | 0.89 | 0.82 | 0.85 |
| **CQ2** | 0.89 | 1.00 | 0.80 | 0.82 | 0.82 | 0.85 | 0.84 | 0.89 |
| **VLB2 A1** | 0.87 | 0.80 | 1.00 | 0.96 | 0.97 | 0.94 | 0.82 | 0.80 |
| **VLB2 D1** | 0.89 | 0.82 | 0.96 | 1.00 | 0.94 | 0.97 | 0.80 | 0.81 |
| **VL20 A1** | 0.87 | 0.82 | 0.97 | 0.94 | 1.00 | 0.96 | 0.84 | 0.81 |
| **VL20 D1** | 0.89 | 0.85 | 0.94 | 0.97 | 0.96 | 1.00 | 0.81 | 0.83 |
| **PD589 A1** | 0.82 | 0.84 | 0.82 | 0.80 | 0.84 | 0.81 | 1.00 | 0.93 |
| **PD589 D3** | 0.85 | 0.89 | 0.80 | 0.81 | 0.81 | 0.83 | 0.93 | 1.00 |

A1 = *V. longisporum* A1 sub-genome, D1 = *V. longisporum* D1 sub-genome and D3 = *V. longisporum* D3 sub-genome. Correlations are calculated with the Spearman’s rank correlation coefficient based on the transcripts per million (tpm) values.

**Table S3. Functional enrichment analysis of *Verticillium longisporum* genes with differential homeolog expression.**

|  |  | **CULTURE MEDIUM** | | | | | | | | | | | **OILSEED RAPE** | | | | | | |
| --- | --- | --- | --- | --- | --- | --- | --- | --- | --- | --- | --- | --- | --- | --- | --- | --- | --- | --- | --- |
|  |  | **VLB2** | | | | **VL20** | | | | **PD589** | | | **VLB2** | | | | **PD589** | | |
| **GO_term** | **Description** | ***p*-value** | **diff expr** | **all** | ***p*-value** | | **diff expr** | **all** | ***p*-value** | | **diff expr** | **all** | ***p*-value** | **diff expr** | **all** | ***p*-value** | | **diff expr** | **all** |
| 0004497 | monooxygenase activity | 1.23E-03 | 9 | 18 | 2.05E-02 | | 7 | 17 |  | |  |  |  |  |  |  | |  |  |
| 0005506 | iron ion binding | 3.48E-03 | 19 | 70 | 3.71E-04 | | 21 | 70 |  | |  |  |  |  |  | 3.34E-02 | | 6 | 51 |
| 0005975 | carbohydrate metabolic process |  |  |  | 2.05E-02 | | 33 | 174 |  | |  |  |  |  |  |  | |  |  |
| 0006725 | cellular aromatic compound metabolic process |  |  |  | 2.32E-02 | | 4 | 6 |  | |  |  |  |  |  |  | |  |  |
| 0008061 | chitin binding |  |  |  | 4.66E-02 | | 5 | 11 |  | |  |  |  |  |  |  | |  |  |
| 0008237 | metallopeptidase activity | 1.34E-02 | 8 | 20 | 3.58E-02 | | 7 | 19 |  | |  |  |  |  |  |  | |  |  |
| 0016020 | membrane |  |  |  |  | |  |  | 4.39E-02 | | 42 | 198 |  |  |  |  | |  |  |
| 0016491 | oxidoreductase activity | 3.38E-05 | 67 | 329 | 1.33E-04 | | 64 | 325 | 3.55E-03 | | 59 | 270 |  |  |  |  | |  |  |
| 0016614 | oxidoreductase activity, acting on CH-OH group of donors | 3.61E-02 | 6 | 14 | 6.47E-03 | | 8 | 18 |  | |  |  |  |  |  |  | |  |  |
| 0016705 | oxidoreductase activity, acting on paired donors, with incorporation or reduction of molecular oxygen | 1.64E-04 | 19 | 56 | 8.56E-05 | | 20 | 57 |  | |  |  |  |  |  | 3.51E-02 | | 5 | 37 |
| 0020037 | heme binding | 9.11E-05 | 23 | 73 | 1.61E-03 | | 20 | 72 |  | |  |  |  |  |  | 1.25E-02 | | 7 | 56 |
| 0022857 | transmembrane transporter activity | 5.97E-09 | 64 | 246 | 2.59E-06 | | 56 | 238 | 3.07E-03 | | 47 | 193 |  |  |  |  | |  |  |
| 0050660 | flavin adenine dinucleotide binding | 1.70E-03 | 21 | 77 | 3.71E-04 | | 23 | 81 |  | |  |  |  |  |  | 4.91E-04 | | 9 | 58 |
| 0055085 | transmembrane transport | 3.38E-05 | 70 | 347 | 2.73E-04 | | 64 | 334 | 3.07E-03 | | 60 | 271 |  |  |  |  | |  |  |
| 0055114 | oxidation-reduction process | 5.97E-09 | 101 | 474 | 2.59E-06 | | 92 | 471 | 3.80E-02 | | 72 | 378 |  |  |  | 3.60E-04 | | 25 | 362 |
| 0071949 | FAD binding | 1.02E-02 | 13 | 43 | 9.16E-03 | | 13 | 43 | 4.39E-02 | | 11 | 31 |  |  |  | 2.79E-04 | | 7 | 26 |
| **COG** | **Description** | ***p*-value** | **diff expr** | **all** | ***p*-value** | | **diff expr** | **all** | ***p*-value** | | **diff expr** | **all** | ***p*-value** | **diff expr** | **all** | ***p*-value** | | **diff expr** | **all** |
| G | Carbohydrate transport and metabolism | 8.39E-05 | 90 | 509 | 5.02E-06 | | 95 | 511 | 2.58E-02 | | 75 | 421 |  |  |  |  | |  |  |
| Q | Secondary metabolites biosynthesis, transport, and catabolism | 1.95E-10 | 73 | 287 | 1.21E-05 | | 59 | 282 | 1.49E-02 | | 49 | 240 |  |  |  |  | |  |  |
| S | Function unknown |  |  |  |  | |  |  | 1.49E-02 | | 249 | 1576 |  |  |  |  | |  |  |
| V | Defence mechanisms |  |  |  |  | |  |  | 1.79E-02 | | 10 | 28 |  |  |  |  | |  |  |

Gene Ontology (GO) terms and Clusters of Orthologous Groups (COGs) that are significantly enriched in genes with differential homeolog expression are displayed. *P*-values were calculated with the Fisher’s exact test and were multiple-testing corrected with the Benjamini-Hochberg method.

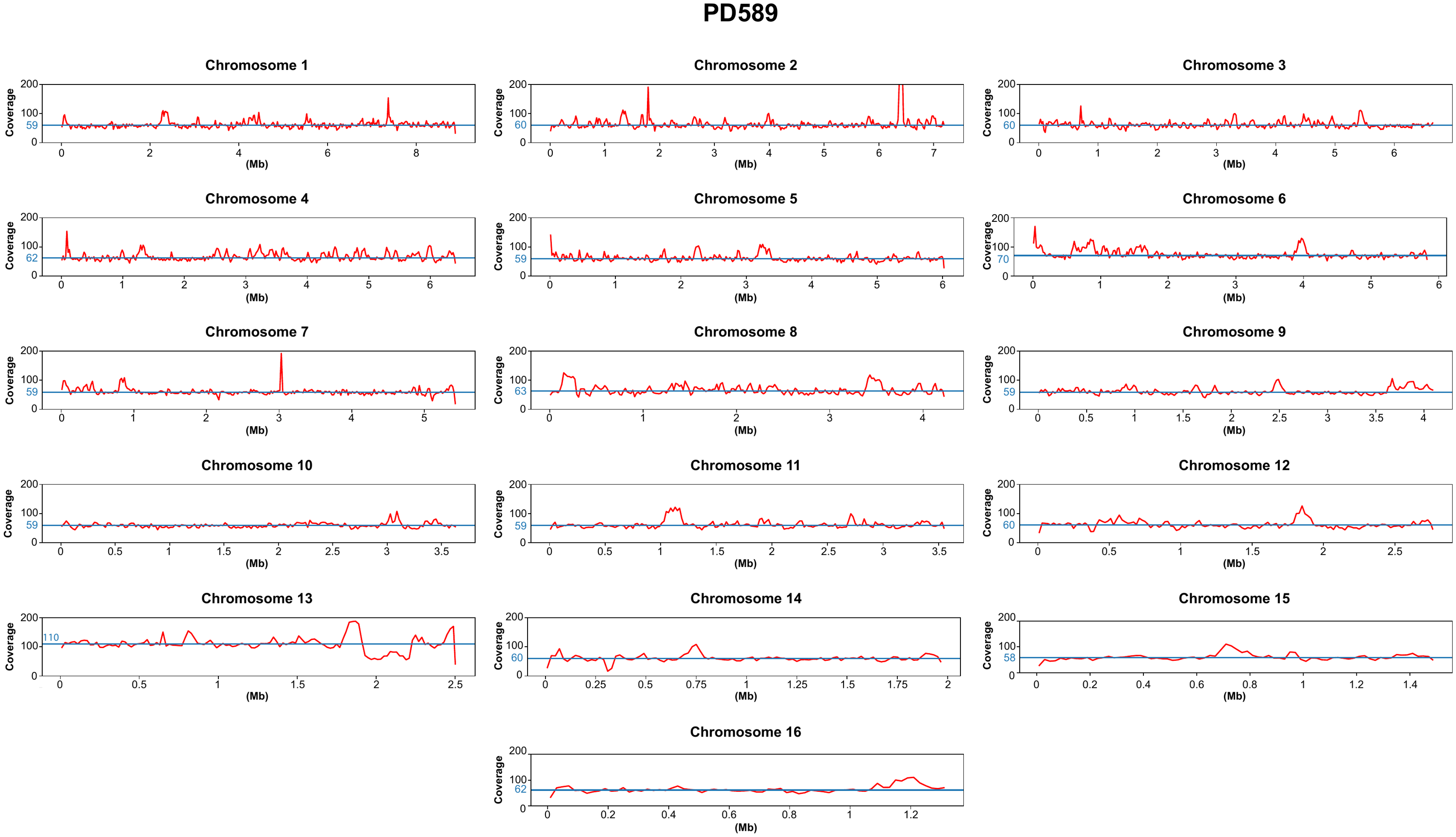

**Figure S1. Read coverage for the *Verticillium longisporum* strain PD589 genome assembly.** The redlines indicates the average coverage of Oxford Nanopore sequenced reads for 20 kb windows. The blue line indicates the median coverage for every individual chromosome.

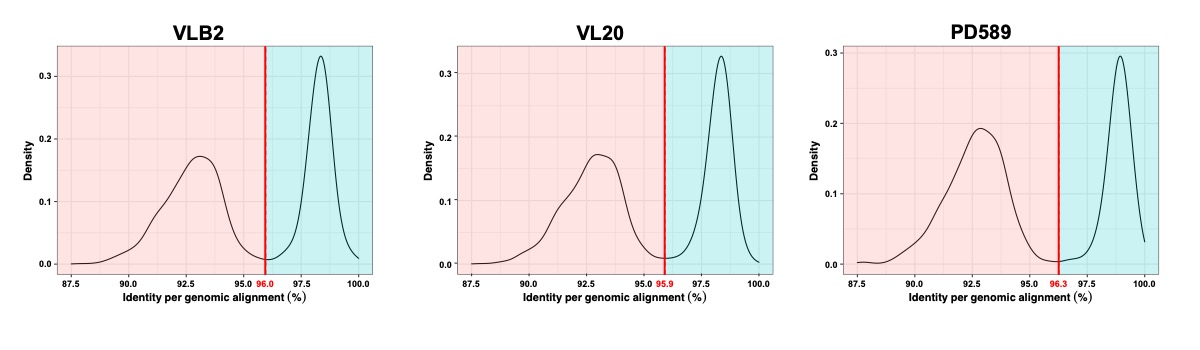

**Figure S2. Density distribution of sequence identity of *Verticillium longisporum* alignments to *Verticillium dahliae*.** *V. longisporum* strains were aligned to *V. dahliae* JR2 with NUCmer. The average sequence identity of these alignments was used to determine smoothed density estimates.

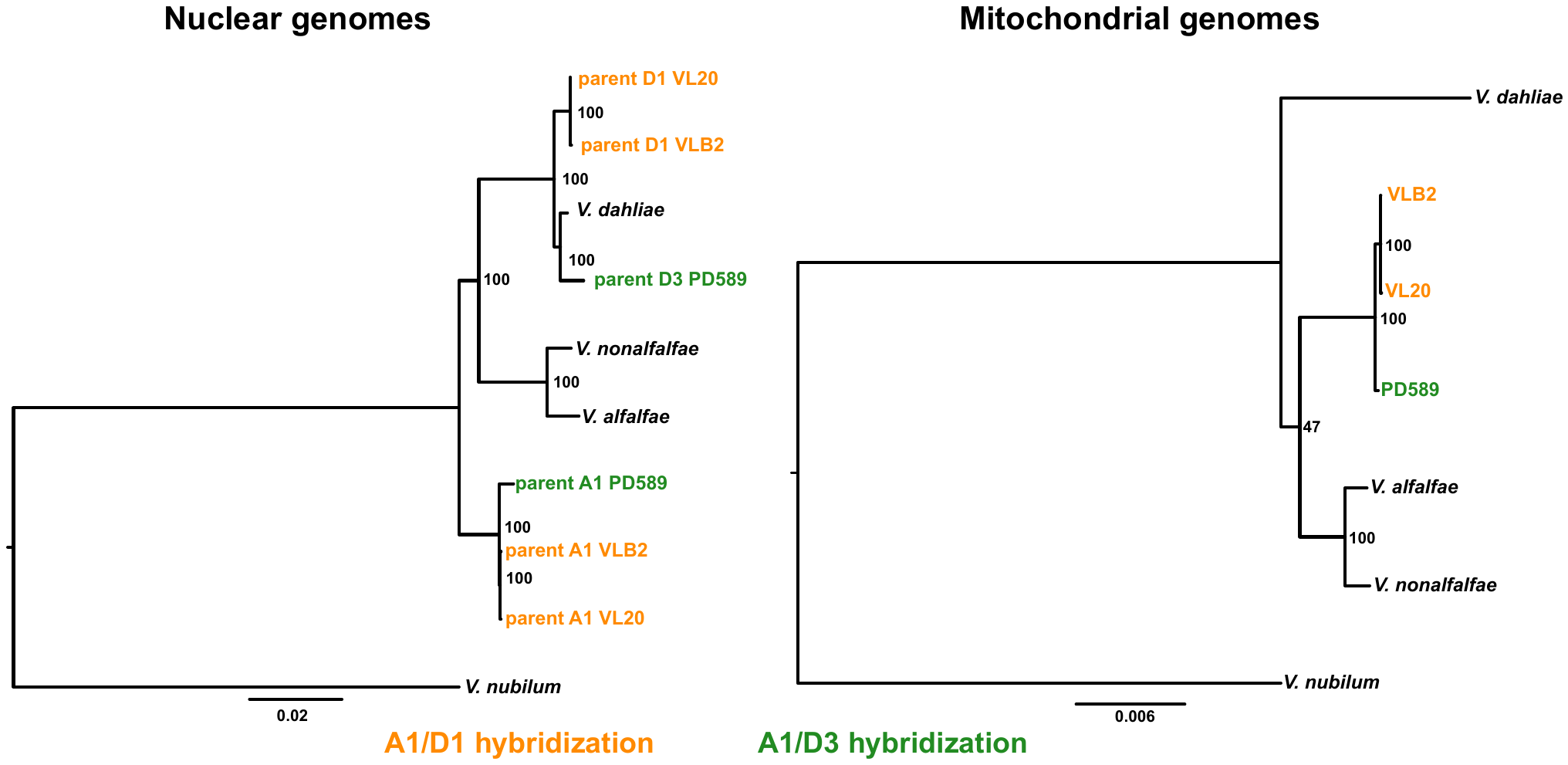

**Figure S3. Phylogenetic relationships between *Verticillium* *longisporum* hybridization parents and haploid *Verticillium* species.** The tree of the nuclear genomes was constructed based on 1,520 Ascomycete benchmarking universal single copy orthologs (BUSCOs) that are present in a single copy in all analyzed *Verticillium* lineages. The tree of the mitochondrial genomes was constructed based on the complete sequence of the mitochondrial genome. The robustness of the inferred phylogeny was assessed by 100 bootstrap approximations.

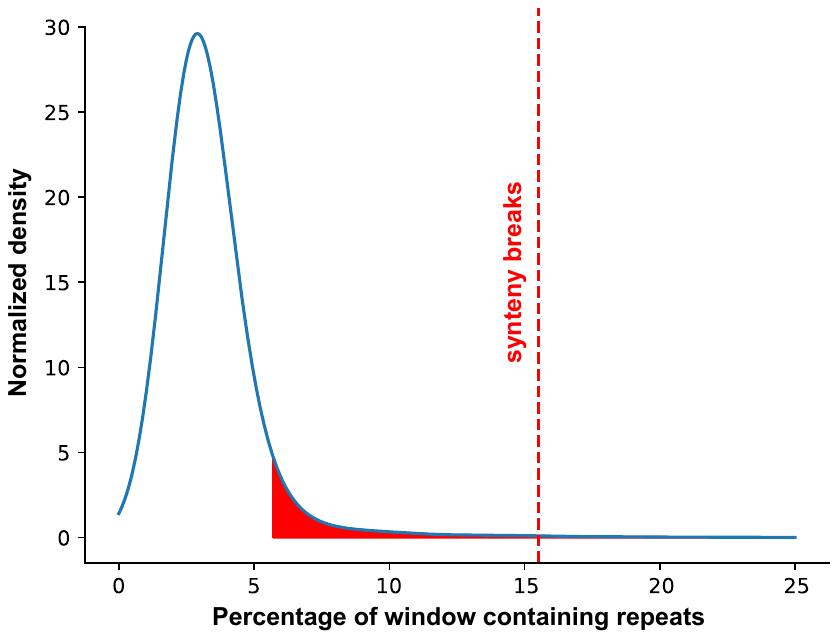

**Figure S4. The association of synteny breaks with repetitive elements.** The black curve represents the normalized density of the median repeat content of 24 randomly chosen 20 kb windows in the *Verticillium longisporum* VL20, which has been permutated 10,000 times. The red line indicates the median repeat content of 20 kb windows around 24 synteny breaks between VLB2 and VL20 (15.5%). The red region indicates the 5% highest data points.
